# Discriminating Lineages of *Batrachochytrium dendrobatidis* using quantitative PCR

**DOI:** 10.1101/2020.08.14.250787

**Authors:** Pria N Ghosh, Ruhan Verster, Thomas Sewell, Simon O’Hanlon, Lola M Brookes, Adrien Rieux, Trenton WJ Garner, Ché Weldon, Matthew C Fisher

## Abstract

The ability to detect and monitor infectious disease in a phylogenetically informative manner is critical for their management. Phylogenetically informative diagnostic tests enable patterns of pathogen introduction or changes in the distribution of genotypes to be measured, enabling research into the ecology of the pathogen. *Batrachochytrium dendrobatidis* (*Bd*), a causative agent of chytridiomycosis in amphibian populations, emerged worldwide in the 21^st^ century and is composed of six lineages which are display varying levels of virulence in their hosts.

Research into the distribution, ecology and pathogenicity of these lineages has been hampered by an inability to type lineage efficiently. Here, we describe a lineage-specific TaqMan qPCR assay that differentiates the two lineages of *Bd* most commonly associated with chytridiomycosis: *Bd*GPL and *Bd*CAPE. We demonstrate how this assay can be used for the surveillance of wild populations of amphibians in Southern Africa using skin swabs, tissue samples and cultured isolates.

## Introduction

*Bd*, a fungal pathogen of amphibians, causes the potentially lethal disease chytridiomycosis. This fungus is a member of the Chytridiomycota, and one of only two species within the phylum known to infect vertebrates (Martel *et al.*, 2013; Longcore & Pessier, 1999). *Bd* has been detected infecting almost 700 species of amphibians, with chytridiomycosis implicated as the cause of population declines in over 500 species (Scheele *et al.*, 2019; Olson & Ronnenberg, 2014). This has led to the extinction of an estimated 90 species of amphibians globally (Scheele *et al.*, 2019).

Within *Bd* there are at least six phylogenetically distinct lineages – *Bd*CAPE, *Bd*GPL, *Bd*ASIA-1, *Bd*ASIA-2/BRAZIL, *Bd*ASIA-3 and *Bd*CH. However, *Bd*CAPE and *Bd*GPL are primarily associated with mass mortality events and species declines or extinctions (Byrne *et al.*, 2019; Farrer *et al.*, 2011; Doddington *et al.*, 2013; Jenkinson *et al.*, 2016; O’Hanlon *et al.*, 2018). While only *Bd*GPL has undergone a rapid, recent and global range expansion, recent research indicates that the other five lineages, which had previously been considered largely spatially restricted, have achieved intercontinental distributions. This raises the possibility of lineage contact and interactions occurring. One such interaction is inter-lineage genetic recombination (Byrne *et al.*, 2019; Jenkinson *et al.*, 2016; O’Hanlon *et al.*, 2018) which may lead to increased virulence (Greenspan *et al.*, 2018). Given the infrequency with which recombinants are isolated, molecular tools that enable the identification and monitoring of areas where they might occur are needed in order to focus research effort effectively. To date, this has been hampered by the lack of a suitable lineage-specific diagnostic test for *Bd*.

Currently, a pan-lineage qPCR assay targeting the ITS region of the genome is the most widely-used method for detecting *Bd* (Boyle *et al.*, 2004). This method has high specificity, is sensitive enough to detect sub-clinical infections, and works with swabs taken from adult amphibian skin or tadpole mouthparts, so is minimally invasive (Garland, Wood & Skerratt, 2011; Kriger *et al.*, 2006; Skerratt *et al.*, 2011; Boyle *et al.*, 2004; Hyatt *et al.*, 2007). A further advantage of this assay working with swab samples is that these are light, easy to carry in difficult field settings and are easy to store (Hyatt *et al.*, 2007; Van Sluys *et al.*, 2008). The primary drawback to this pan-lineage qPCR assay is its inability to type the lineage of *Bd* present in a sample. Although there have been attempts to design lineage-specific diagnostics based on the targeted genomic region, the ITS is unable to resolve inter-specific phylogenies within *Bd* owing to a breakdown in concerted evolution across this multi-allelic array (O’Hanlon *et al.*, 2018; Schoch *et al.*, 2012).

The most accurate method for determining lineage is whole genome sequencing (WGS) from a pure culture of *Bd*. Although comparative genomic analyses have been extremely valuable in the field of chytrid research, the expense and requirement for a pure isolate culture make this approach impractical on the large scale required to answer fundamental questions about *Bd* lineage distribution and ecology. The most reliable alternative approach to WGS, multilocus sequence typing (MLST) is suitable for use with swabs but requires access to a specialised technology, the Fluidigm Access Array platform, and a reasonably high infection burden (>150 genomic equivalents (GE)). It is, therefore, not applicable for all *Bd* positive cases, especially in hosts with a subclinical infection burden (Byrne *et al.*, 2017).

Recent research has shown that phylogenetic analyses based on *Bd* mitochondrial DNA (mtDNA) recovers the same lineages as those recovered by WGS (O’Hanlon *et al.*, 2018). The *Bd* mitochondrial genome has recently been fully sequenced, has a low rate of recombination relative the nuclear genome, and is present at high copy number so is likely to be detectable at low infection burdens. There is also a precedent for using mtDNA for phylogenetic studies across a wide range of taxa (Barr *et al.*, 2011; Li *et al.*, 2016; Penry *et al.*, 2018; Schreeg *et al.*, 2016; Song *et al.*, 2016). Here, we report a lineage-specific qPCR assay for *Bd* based on mtDNA which detects the two lineages of *Bd* most commonly associated with chytridiomycosis in wild populations – *Bd*GPL and *Bd*CAPE – with high specificity and sensitivity.

## Materials and Methods

### Assay design and optimisation

A 175,295 kb long mtDNA alignment for 150 chytrid isolates, with 145 isolates of *Bd* representing *Bd*GPL, *Bd*CAPE, *Bd*ASIA-1, *Bd*ASIA-2/BRAZIL and *Bd*CH, was manually screened for lineage-specific single nucleotide polymorphisms (SNPs) (O’Hanlon *et al.*, 2018). Candidate primers and probes were designed using Primer Express Software 3.0 (ThermoFisher Scientific, Massachusetts, USA).

We optimised primer and probe concentrations separately using a checkboard system in singleplex with DNA quantitation standards for the DNA template. We selected final primer and probe concentrations which maximised reaction efficiency compared to oligonucleotide concentration, while ensuring that the mean Ct values for a given quantity of DNA remained as similar as possible between the *Bd*CAPE and *Bd*GPL reactions. Consistency between the reactions made it possible to multiplex the two qPCR reactions into a single assay diagnosing both *Bd*GPL and *Bd*CAPE. All optimisation and validation reactions were carried out on both a 7300 Real-Time PCR system and a QuantStudio 7 Flex Real-Time PCR system (Applied Biosystems, Foster City, CA, USA).

The cycling conditions were identical to Boyle *et al*’s pan-lineage *Bd* diagnostic, except that we reduced the number of cycles from 50 to 40 and increased the annealing temperature from 60°C to 62°C in order to increase the specificity. We set the Ct threshold for all reactions at ΔRn = 0.1, and the total reaction volume in all cases was 25μl in total (20μl master mix and 5μl DNA template). The optimised multiplex reaction mix and full cycling conditions are provided in the supplementary information. When the reactions were run in singleplex the same quantities of all reactions were used, except for an increase in the volume of filtered water commensurate with the quantity of primer and probe removed. We quantified the starting DNA present in a reaction using standard curve analysis calculated from DNA quantitation standards made from isolate IA042 (*Bd*GPL) and isolate TF5a1 (*Bd*CAPE) (supplementary information). Both these isolates were previously typed to their respective lineages using WGS (O’Hanlon *et al.*, 2018). The final primer and TaqMan MGB probe sequences are shown in Figure 1. We carried out pan-lineage *Bd* detection using Boyle *et al*’s protocol (Boyle *et al.*, 2004).

**Figure 1:**
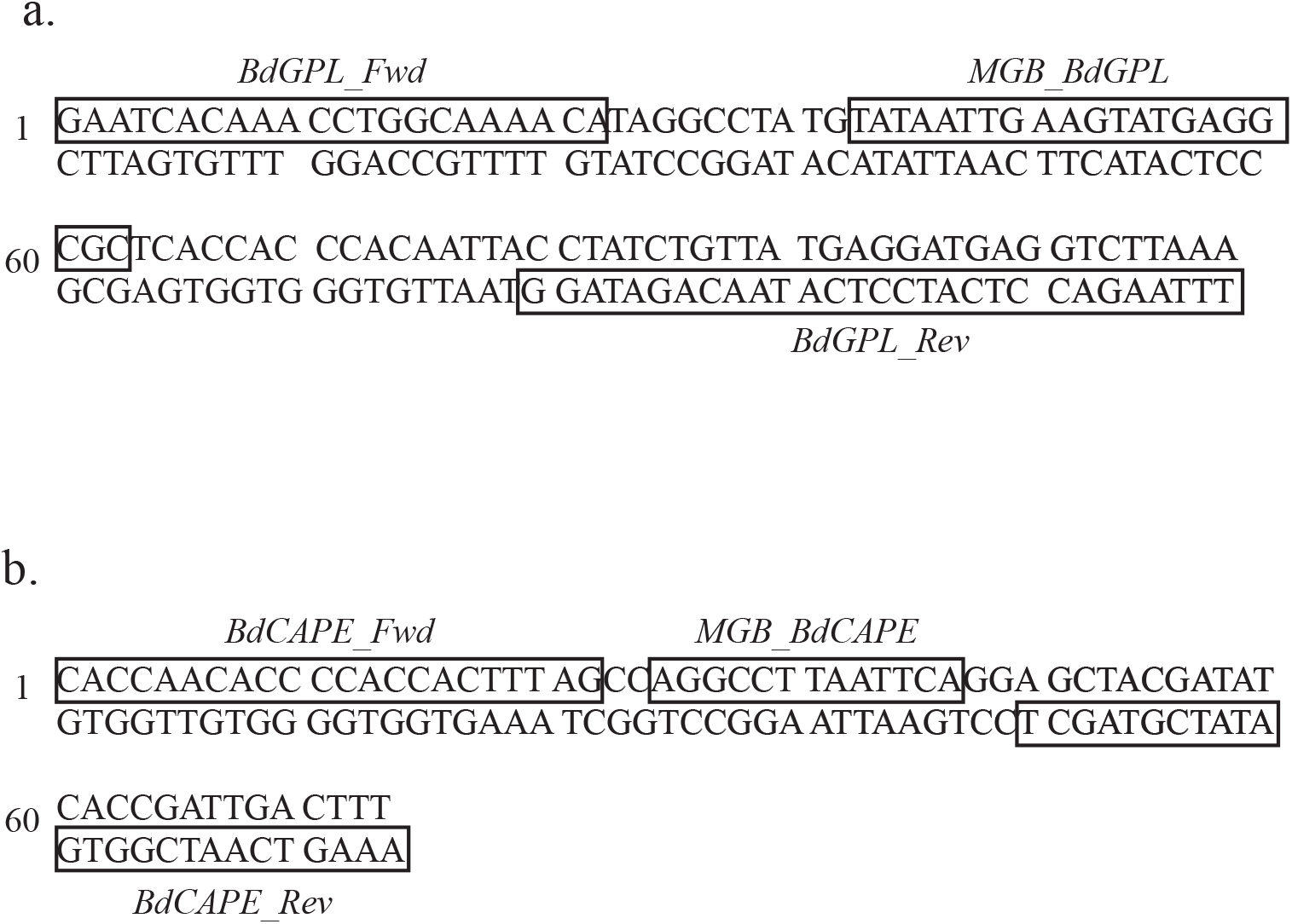
*Bd* mtDNA sequence sections showing the a) *Bd*CAPE-specific and b) *Bd*GPL-specific primer and TaqMan MGB probe sequences.

### Assessing lineage-specific qPCR specificity, sensitivity and reliability

A panel of 52 isolates (38 *Bd*GPL, six *Bd*CAPE, four *Bd*ASIA-1, three *Bd*ASIA-2/BRAZIL and one *Bd*CH) collected from 17 countries as well as from traded amphibians was used to test the specificity of the lineage-specific qPCR (Figure 2a). All isolates had been previously assigned a lineage by WGS (O’Hanlon *et al.*, 2018) (Figure 2b). We extracted DNA from the live cultures following RACE protocols (Supplementary information) for 50 isolates. DNA extracts for isolates UM142 and CLFT-065 were provided by University of Michigan. We tested all 52 isolates with both the lineage-specific qPCR reactions in singleplex to check for cross-reactivity. All other reactions were carried out in multiplex. To test for sensitivity, we ran the lineage-specific qPCR assay with a 10-fold dilution series of *Bd*GPL and *Bd*CAPE DNA quantitation standards in duplicate. Once a suitable range of DNA concentrations for standard curve analysis had been identified, we tested the repeatability of the assay within and between plates by comparing reaction efficiency, and R^2^ and Ct values.

**Figure 2:**
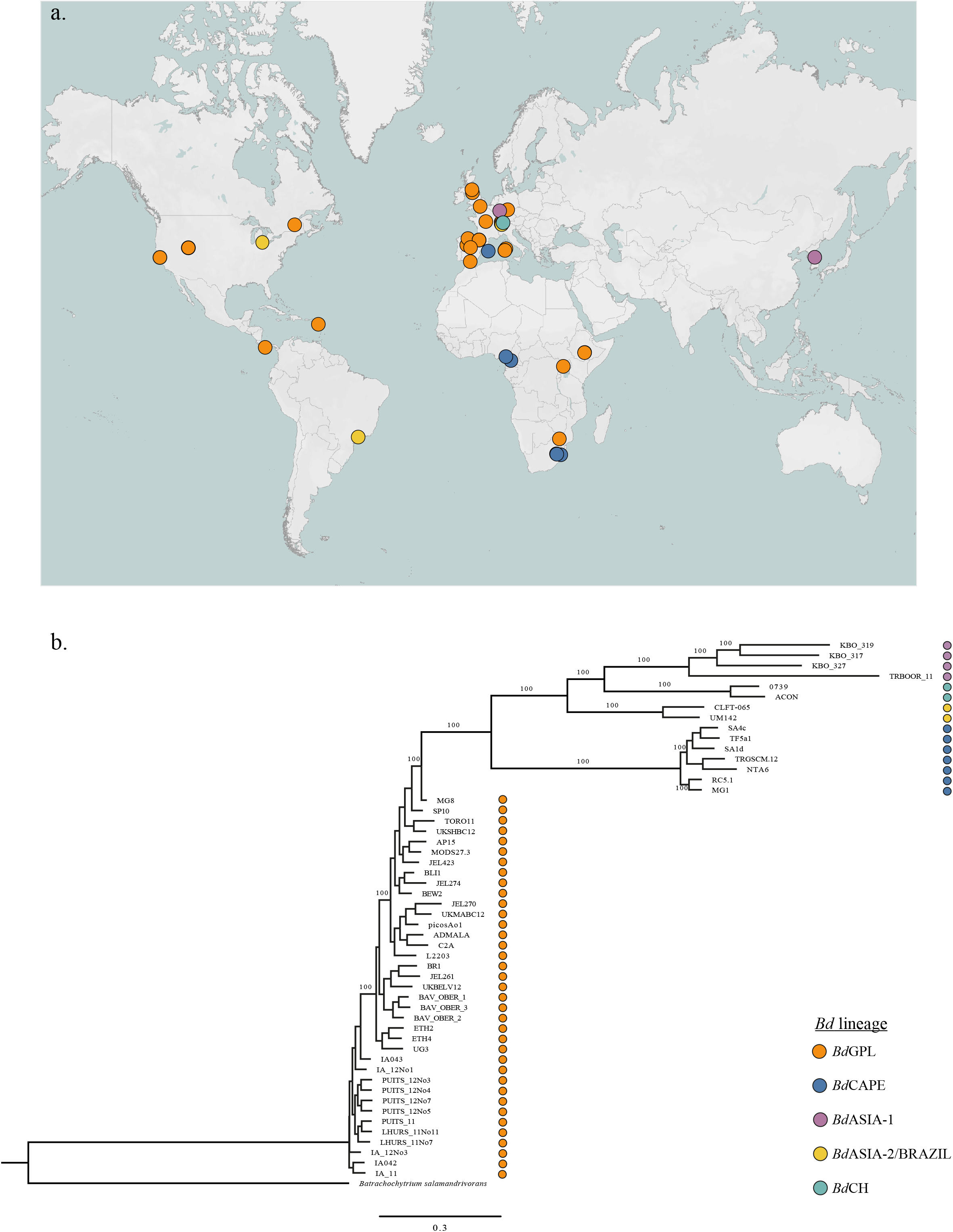
a) Map showing source locations of *Bd* isolates used for assay specificity testing; b) phylogeny of isolates used for assay specificity testing, generated from WGS analysis. Lineage is indicated by dot colour. Map generated in Tableau^TM^ and formatted in Adobe Illustrator^®^.

### Field sample collection and processing

WGS analysis has revealed that *Bd*GPL and *Bd*CAPE are both present in wild populations in South Africa in the Orange River Basin (Figure 3a). This region was therefore selected as a study system to see if the lineage-specific qPCR assay diagnosed the same lineages from field sites as WGS. Sites were surveyed along the Orange River at approximately 100 km apart from the source in the Drakensburg Mountains to near the estuary on the border with Namibia. At each site, we attempted to culture and isolate *Bd* using non-lethal tissue sampling from adult amphibians and lethal mouthpart sampling from tadpoles (Fisher *et al.*, 2018). Any resulting chytrids were lineage-typed by WGS using the protocol described by O’Hanlon *et al* (O’Hanlon *et al.*, 2018). All adult amphibians were also swabbed following ‘*Risk Assessment of Chytridiomycosis to European Amphibian biodiversity* (*RACE*)’ protocols (Biodiversa, 2013; Fisher *et al.*, 2018). DNA extraction also followed the *RACE* protocol (supplementary information), following which swab samples were tested for *Bd* using the pan-lineage qPCR diagnostic. Any samples which were positive in duplicate with an infection load of more than one genomic equivalent (GE) were tested with the lineage-specific qPCR diagnostic in duplicate with DNA quantitation standards and two no-template controls.

**Figure 3:**
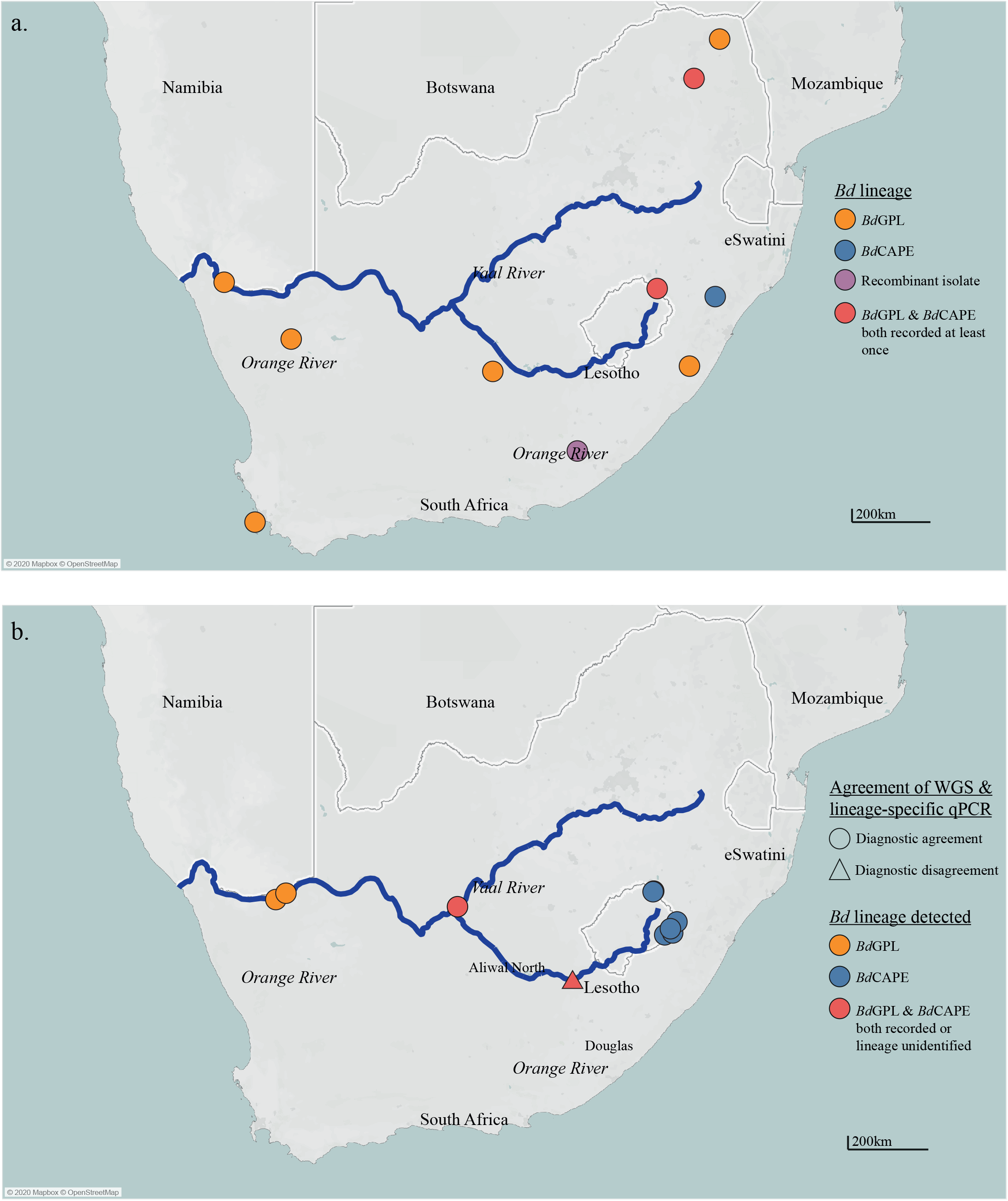
Maps showing a) *Bd* lineage locations in South Africa identified by isolation and whole genome sequence analysis and b) comparison of *Bd* lineage typing of sites in South Africa using WGS analysis and lineage-specific qPCR. Agreement of diagnostics is indicated by shape of the mark; lineages identified at the site are identified by the colour. Map generated in Tableau™ and formatted in Adobe Illustrator^®^, additional data downloaded from naturalearthdata.com.

## Results

### Lineage-specific qPCR performance under laboratory conditions

The *Bd* lineage-specific assay was able to detect all isolates assigned to the relevant lineage by WGS. The *Bd*GPL assay also detected DNA from one isolate from the *Bd*ASIA-2/BRAZIL lineage (UM142) but at an unrealistic concentration. This reaction also did not cross the reaction threshold during the exponential phase, as would be expected in a true positive result (Figure S1). Both lineage-specific assays had a limit of detection of 1 GE (Figure 4). The *Bd*CAPE-specific qPCR recorded a plateauing of fluorescence at 10,000 GE, resulting in an underestimation of *Bd* load when comparing with the other quantitation standard fluorescence levels (Figure 4a). This plateauing owes to too much target DNA present in a sample quenching the PCR reaction due to the depletion of reagents. Based on these results, we recommend a DNA quantitation standard range for both lineage-specific qPCRs of 1000 GE, 100 GE, 10 GE and 1 GE for standard curve analysis.

**Figure 4:**
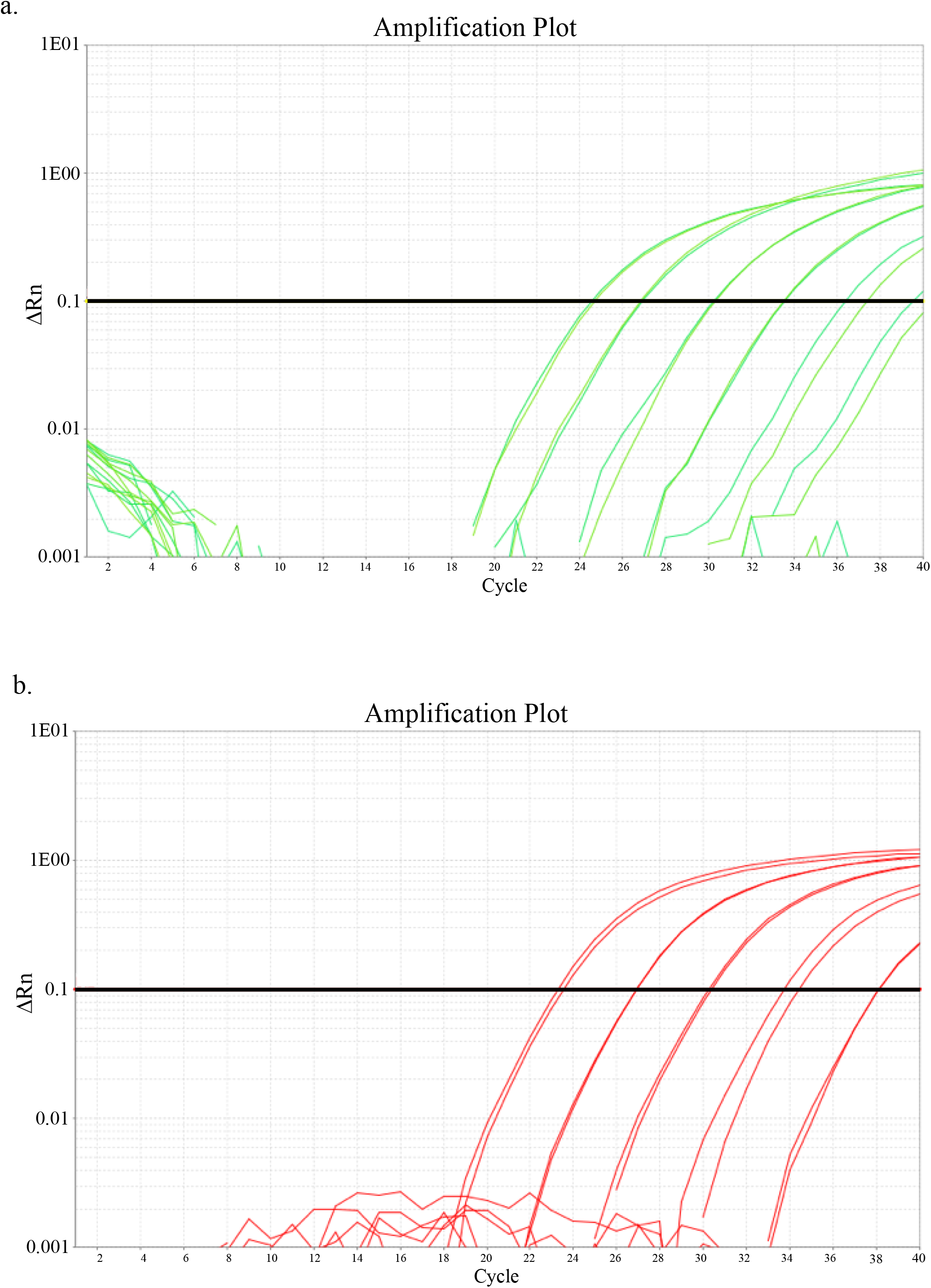
qPCR fluorescence amplification plots showing amplification of 10,000GE, 1000GE, 100GE, 10GE and 1GE DNA quantitation standards with a) *Bd*CAPE-specific primers and TaqMan MGB probe and b) *Bd*GPL-specific primers and TaqMan MGB probe.

Both assays performed well over three separate qPCR plates. The overall mean percentage efficiency for the *Bd*CAPE-specific qPCR was 108.87% (ranging from 99.35% to 126.32%), and for the *Bd*GPL-specific qPCR was 97.35% (ranging from 92.93% to 104.21%). All R^2^ values, whether averaged across and within plates were >0.95 (ranging from 0.98 to 0.99) and the Ct values of DNA quantitation standards were also consistent, all crossing the Ct threshold within one cycle of each other in all cases.

### Lineage-specific qPCR performance under field conditions

We obtained at least one pure culture of *Bd* and at least one *Bd*-positive skin swab that met the threshold for qPCR lineage-testing from 10 sampling sites. At all sites except at Aliwal North, WGS analysis and the lineage-specific qPCR agreed in their lineage typing (Figure 3b). At Aliwal North, the lineage-specific qPCR was positive for *Bd*CAPE, but WGS analysis revealed that the isolate recovered from the site was a CAPE/GPL recombinant. At Douglas, a single swab returned a positive result for both *Bd*GPL and *Bd*CAPE. Both lineages were also recovered from the same pond at Douglas by isolation and WGS.

## Discussion

We have designed a *Bd*CAPE- and *Bd*GPL-specific qPCR diagnostic that will facilitate research into the ecology and distributions of these epidemiologically important *Bd* lineages. The diagnostic gap that has persisted until now resulted in an inability to gather baseline data on *Bd* distribution at a phylogenetically and spatially useful scale. This lack has led to difficulties investigating the ecology and epidemiological significance of lineage interactions. Our lineage-specific qPCR complements existing diagnostics for *Bd* while meeting the need to decipher lineage in large numbers of samples and in animals with low infection burdens.

The qPCR assay was able to correctly detect whether *Bd*GPL or *Bd*CAPE was present in 51 out of 52 isolates. The most likely explanation for the low-level cross-reactivity of the *Bd*GPL-specific qPCR assay with a *Bd*ASIA-2/BRAZIL isolate is that the DNA extract was contaminated with DNA from a *Bd*GPL isolate at some point after WGS had been carried out. Sequencing reads from this isolate’s mtDNA were screened for the presence of the *Bd*GPL-specific probe, which was not found. This result highlights the importance of manually examining amplification plots to assess whether they appear atypical, and to continue to utilise WGS methods where possible to corroborate results from the lineage-specific qPCR.

The assay was sufficiently sensitive for use in cases of low infection burden, with a sensitivity of one genomic equivalent. We note however that mtDNA copy number has not been quantified for any *Bd* isolate and that ITS copy number, which is used to quantify infection burden in the majority of *Bd* surveillance studies, is known to be highly variable (Rebollar *et al.*, 2017; Longo *et al.*, 2013). We recommend therefore that in order to remain consistent with the current literature, that infection intensity is reported based on the pan-lineage *Bd* diagnostic, but that this quantification should be viewed as semi-quantitative rather than absolute. Under experimental conditions, this problem could be resolved by quantifying the copy number of the targeted region for specific isolates before carrying out any experimental work.

Novel diagnostics are challenging to test in a field setting, due to the lack of *a priori* knowledge of the location of the target pathogen. However, agreement of the lineage-specific qPCR diagnostic with WGS analysis for sites in South Africa strongly supports the use of the qPCR assay under field conditions. Crucially, the results from Douglas indicate that the assay is able to identify where mixed-lineage infections at both the population and individual host level are occurring. This indicates that the assay is appropriate for use in putative lineage contact zones where lineage interactions may occur. The lineage-specific qPCR identified the only recombinant lineage collected as *Bd*CAPE. This suggests uniparental inheritance of mitochondria in *Bd*, a new insight into the pathogen’s biology.

The consistency of any diagnostic assay across multiple laboratories is essential to its utility and ease of use. The wide application of the pan-lineage *Bd* qPCR diagnostic to date means that any laboratory working on *Bd* is likely to have access to all the reagents and facilities required to perform the lineage-specific assay described here, at a similar cost per sample. We recommend that similar lineage-specific mtDNA based diagnostics for *Bd*ASIA-1, *Bd*ASIA-2/BRAZIL, *Bd*ASIA-3 and *Bd*CH are designed as a research priority. This would allow the rapid and widespread generation of baseline data from field samples, allowing much finer delineation of *Bd* lineage distributions globally and further insight into *Bd* ecology. Lineage-specific diagnostics also enable *in vivo* and *in vitro* experimental manipulation of *Bd* lineages, thus providing further insight into *Bd* population dynamics and structure.

## Acknowledgements

PNG and MCF were supported by the Morris Animal Foundation, the Leverhulme Trust (RPG-2014-273), the Natural Environment Research Council (NERC NE/E006841/1) and a PhD DTP award by the ICL Grantham Institute. MCF is a fellow in the CIFAR ‘Fungal Kingdoms’ program. The authors would like to thank Tim James, University of Michigan for providing DNA extract of isolates UM42 and CLFT-065.

## Data Accessibility

The mtDNA alignment used to design the primers and probes will be uploaded to Dryad open repository.

## Author contributions

The research was planned by PNG, TWJG & MCF. mtDNA sequences were compiled and aligned by AR. Primer and TaqMan MGB probe design was done by PNG. Assay validation was carried out by PNG and LMB. PNG, RV, TWJG and CW carried out fieldwork in South Africa. SOH and TS carried out whole genome sequences analyses. PNG wrote the article with input from all authors.

