## Supplementary methods and Figure S1 for "Discriminating Lineages of *Batrachochytrium dendrobatidis* using quantitative PCR"

Supplementary materials for Discriminating Lineages of *Batrachochytrium dendrobatidis* using quantitative PCR

### Materials and Methods

#### Bd *lineage-specific qPCR reaction mix and cycling conditions*

The optimised multiplex reaction mix for a single 25µl reaction (with a starting concentration of 100µM for all primers and probes) was as follows: 12.5µl TaqMan Fast Advanced MasterMix (ThermoFisher Scientific, Massachusetts, USA); 0.125µl each of *Bd*GPL-specific forward and reverse primers; 0.225µl each of *Bd*-CAPE specific forward and reverse primers; 0.03125µl *Bd*GPL-specific TaqMan MGB probe; 0.09375µl of *Bd*-CAPE specific TaqMan MGB probe; 0.2µl Bovine Serum Albumin (BSA); 6.475µl filtered water.
The cycling conditions were two minutes at 50°C, followed by 10 minutes at 95°C, then 15 seconds at 95°C and one minute at 62°C, cycled 40 times. The Ct threshold was set at ΔRn = 0.1 for all reactions.

#### Risk Assessment of Chytridiomycosis to European Amphibian biodiversity *(*RACE*) swabbing protocol*

Use tubed sterile dry swabs with a fine tip (if possible). Ideally use gloves when handling amphibians and change gloves between each animal. Alternatively, thoroughly wash/disinfect hands between each animal however this will not prevent cross-contamination of samples with DNA. Hold the animal firmly by its back legs. For caudatans, hold the animal on either side of its abdomen. Firmly run the swab over the back and legs ten times. You will need to press quite hard on the skin as you swab, as you are trying to remove any fungal zoospores or sporangia embedded on the amphibian's skin. Rotate the swab to ensure the entire surface has come into contact with the skin. Next, swab the back legs and toes ten times per limb, then flip the animal over to swab the pelvic patch and insides of the back legs ten tens. Pay particular attention to the swabbing the back legs, toes and pelvic patch, because this is where *Bd* is most likely to be found. Release the animal and store the swab at four degrees. Ambient temperature is fine for a few hours. Give the swab a clear number either in permanent marker or pencil. In a separate data file record for each swab: location (as specific as possible); species (if known); collector's name; date; any additional information that may be relevant e.g. male/female/juvenile/body condition).

#### RACE *DNA extraction protocol*

In a BioSafety ACDP Class 2 cabinet or equivalent, fill one 1.5ml Eppendorf tube with approximately 0.03g Zirconium/silica microbeads for each sample. Pipette 60µl of Prepman ultra (ThermoFisher Scientific, Massachusetts, USA) into each Eppendorf for swab samples and 50µl of Prepman ultra for tissue samples. If processing swab samples, snap the end of each swab off into an individual Eppendorf tube directly. If processing tissue samples, add approximately 1-4$mm^2$ of tissue, depending on the thickness. Large tissue samples should be dissected for processing. Add the tissue to an Eppendorf tube. Homogenize the samples using a bead beater for 45 seconds, the centrifuge for 30 seconds at 14500rpm. Repeat the homogenisation and centrifugation. Incubate the samples at 100°C for 10 minutes, followed by 2 minutes cooling at ambient temperature. Centrifuge for 3 minutes at 14500 rpm. Collect as much supernatant as possible and store in a fresh sterile 0.5ml Eppendorf tube.

*Preparation of DNA quantitation standards*

All work with live *Bd* isolate cultures should be carried out in a BioSafety ACDP Class 2 cabinet or equivalent. To prepare DNA quantitation standards, transfer 500ml of *Bd* culture suspended in Tryptone Gelatin hydrosolate Lacttose (TGhL) broth to a TGhL 1% agar plate. Allow the plate to dry, the para-film it and incubate inverted at 18°C for approximately 5 days. When large numbers of zoospores have been produced, wash the plates with 1ml of TGhL broth and incubate them for 10 minutes at ambient temperature. Pipette the broth off the plate and collect the supernatant. Repeat this process at least twice. Calculate the concentration of live zoospores in the supernatant using a haemocytometer, then dilute down to the appropriate concentration of zoospores with more TGhL broth. Transfer 1ml of the solution into a 1.5ml sterile Eppdendorf tube and centrifuge for 10 minutes at 14500 rpm. Remove the supernatant, being careful not to disturb the pellet, and then add 50μl of Prepman Ultra. After this point, follow the RACE protocol for DNA extraction.

*Table S1. Isolates used for DNA quantitation standards*

| Isolate | Lineage | Source location | Latitude | Longitude | Year isolated | Host species |
|---|---|---|---|---|---|---|
| TF5a1 | *Bd*CAPE | Mallorca | 39.86 | -2.84 | 2007 | *Alytes muletensis* |
| IA042 | *Bd*GPL | Spain | 40.85 | -3.96 | 2004 | *Alytes obstetricans* |

*Table S2. Isolates used for lineage-specific qPCR specificity testing panel*

| Isolate | Lineage | Source location | Year isolated | Host species |
|---|---|---|---|---|
| KBO_319 | *Bd*ASIA-1 | South Korea | 2014 | *Bombina orientalis* |
| KBO_317 | *Bd*ASIA-1 | South Korea | 2014 | *Bombina orientalis* |
| KBO_327 | *Bd*ASIA-1 | South Korea | 2014 | *Bombina orientalis* |
| TRBOOR_11 | *Bd*ASIA-1 | Trade | 2011 | *Bombina variegata* |
| 0739 | *Bd*CH | Switzerland | 2007 | *Alytes obstetricans* |
| ACON | *Bd*CH | Switzerland | 2007 | *Alytes obstetricans* |
| CLFT-065 | *Bd*ASIA2/BRAZIL | Brazil | 2013 | *Hylodes japi* |
| UM142 | *Bd*ASIA2/BRAZIL | Brazil | 2009 | *Lithobates catesbeianus* |
| SA4c | *Bd*CAPE | South Africa | 2010 | *Amietia delalandii* |
| TF5a1 | *Bd*CAPE | Mallorca | 2007 | *Alytes muletensis* |
| SA1d | *Bd*CAPE | South Africa | 2010 | *Hadramophryne natalensis* |
| TRGSCM.12 | *Bd*CAPE | Trade | 2007 | *Geotrypetes seraphini* |
| NTA6 | *Bd*CAPE | Cameroon | 2012 | *Ophisthothylax immaculatus* |
| RC5.1 | *Bd*CAPE | France | 2010 | *Lithobates catesbeianus* |
| MG1 | *Bd*CAPE | South Africa | 2008 | *Amietia hymenopus* |
| MG8 | *Bd*GPL | South Africa | 2008 | *Amietia delalandii* |
| SP10 | *Bd*GPL | Sardinia | 2010 | *Discoglossus sardus* |
| TORO11 | *Bd*GPL | Spain | 2011 | *Alytes obstetricans* |
| UKSHBC12 | *Bd*GPL | UK | 2012 | *Bufo calamita* |
| AP15 | *Bd*GPL | Sardinia | 2010 | *Discoglossus sardus* |
| MODS27.3 | *Bd*GPL | Sardinia | 2010 | *Discoglossus sardus* |
| JEL423 | *Bd*GPL | Panama | 2004 | *Hylomantus lemur* |
| BLI1 | *Bd*GPL | Switzerland | 2010 | *Alytes obstetricans* |
| JEL274 | *Bd*GPL | USA | 1999 | *Bufo boreas* |
| BEW2 | *Bd*GPL | Switzerland | 2010 | *Alytes obstetricans* |
| JEL270 | *Bd*GPL | USA | 1999 | *Lithobates catesbeianus* |
| UKMABC12 | *Bd*GPL | UK | 2012 | *Bufo calamita* |
| picosAo1 | *Bd*GPL | Spain | 2010 | *Alytes obstetricans* |
| ADMALA | *Bd*GPL | Spain | 2012 | *Alytes dickilleni* |
| C2A | *Bd*GPL | Spain | 2002 | *Alytes obstetricans* |
| L2203 | *Bd*GPL | Montserrat | 2009 | *Leptodactylus fallax* |
| BR1 | *Bd*GPL | Switzerland | 2011 | *Alytes obstetricans* |
| JEL261 | *Bd*GPL | Canada | 1999 | *Lithobates catesbeianus* |
| UKBELV12 | *Bd*GPL | UK | 2012 | *Lissotriton vulgaris* |
| BAV_OBER_1 | *Bd*GPL | Germany | 2012 | *Alytes obstetricans* |
| BAV_OBER_2 | *Bd*GPL | Germany | 2013 | *Alytes obstetricans* |

| | | | | |
|---|---|---|---|---|
| BAV_OBER_3 | *Bd*GPL | Germany | 2013 | *Alytes obstetricans* |
| ETH2 | *Bd*GPL | Ethiopia | 2011 | *Leptopelis spp.* |
| ETH4 | *Bd*GPL | Ethiopia | 2011 | *Afrixalus enseticola* |
| UG3 | *Bd*GPL | Uganda | 2012 | *Amietophrynus spp.* |
| IA043 | *Bd*GPL | Spain | 2004 | *Alytes obstetricans* |
| IA_12No1 | *Bd*GPL | Spain | 2012 | *Alytes obstetricans* |
| PUITS_12No3 | *Bd*GPL | France | 2012 | *Alytes obstetricans* |
| Puits_12No4 | *Bd*GPL | France | 2012 | *Alytes obstetricans* |
| Puits_12No5 | *Bd*GPL | France | 2012 | *Alytes obstetricans* |
| Puit_12No7 | *Bd*GPL | France | 2012 | *Alytes obstetricans* |
| PUITS_11 | *Bd*GPL | France | 2011 | *Alytes obstetricans* |
| LHURS_11No11 | *Bd*GPL | France | 2011 | *Alytes obstetricans* |
| LHURS_11No7 | *Bd*GPL | France | 2011 | *Alytes obstetricans* |
| IA_12No3 | *Bd*GPL | Spain | 2012 | *Alytes obstetricans* |
| IA042 | *Bd*GPL | Spain | 2004 | *Alytes obstetricans* |
| IA_11 | *Bd*GPL | Spain | 2011 | *Alytes obstetricans* |

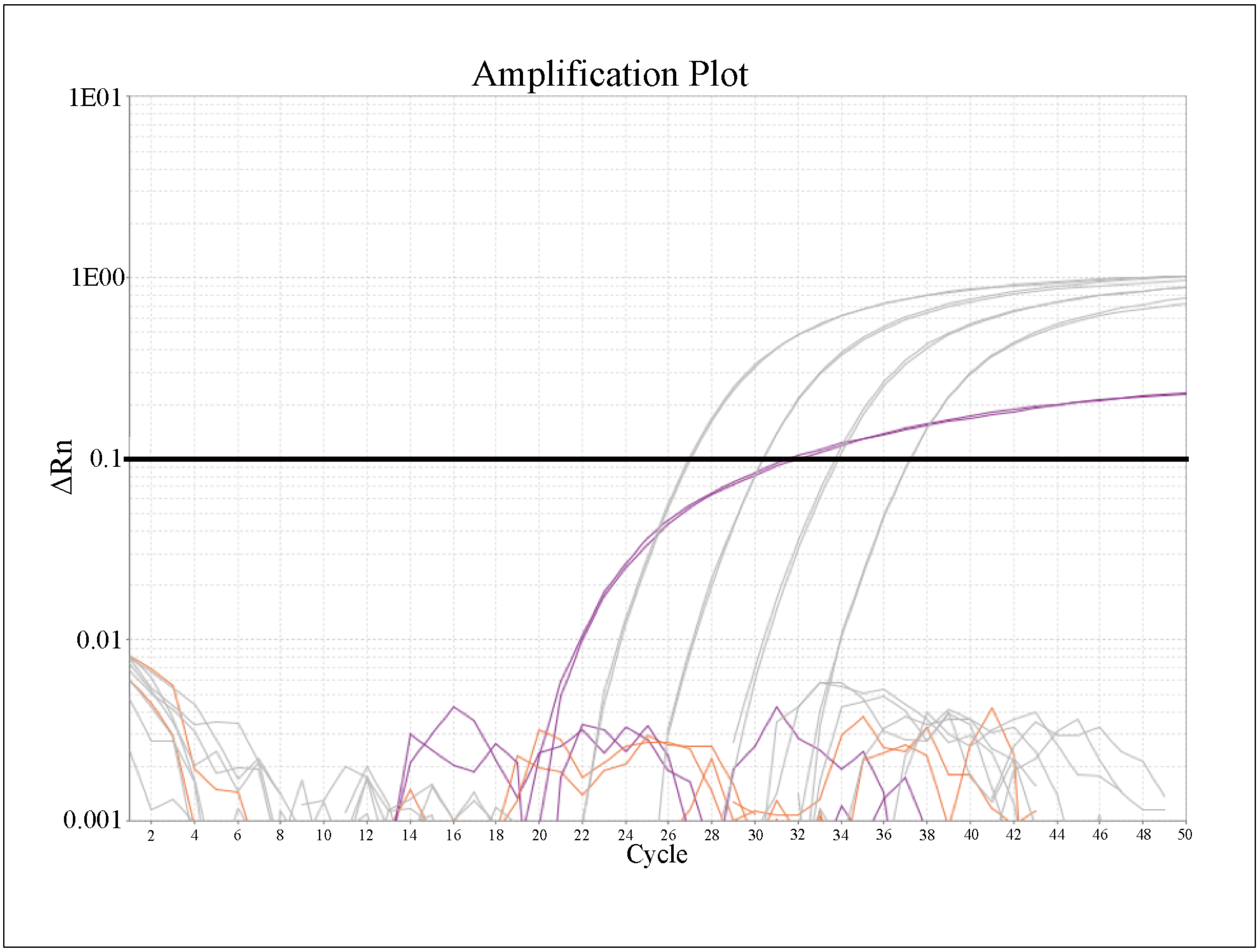


*Figure S1: qPCR fluorescence amplification plot for isolate UM142 (purple) with* Bd*GPL-specific primers and probes and DNA quantitation standards for* Bd*GPL of 1000GE, 100GE, 10GE and 1GE (grey).*
